# High-Efficiency Electroporation of Chytrid Fungi

**DOI:** 10.1101/2020.05.25.114942

**Authors:** Andrew J.M. Swafford, Shane P. Hussey, Lillian K. Fritz-Laylin

## Abstract

Two species of parasitic fungi from the phylum Chytridiomycota (chytrids) are annihilating global amphibian populations. These chytrid species—*Batrachochytrium dendrobatidis* and *B. salamandrivorans*—have high rates of mortality and transmission. Upon establishing infection in amphibians, chytrids rapidly multiply within the skin and disrupt their hosts’ vital homeostasis mechanisms. Current disease models suggest that chytrid fungi locate and infect their hosts during a motile, unicellular ‘zoospore’ life stage. Moreover, other chytrid species parasitize organisms from across the tree of life, making future epidemics in new hosts a likely possibility. Efforts to mitigate the damage and spread of chytrid disease have been stymied by the lack of knowledge about basic chytrid biology and tools with which to test molecular hypotheses about disease mechanisms. To overcome this bottleneck, we have developed high-efficiency delivery of molecular payloads into chytrid zoospores using electroporation. Our electroporation protocols result in payload delivery to between 75-97% of living cells of three species: *B. dendrobatidis, B. salamandrivorans,* and a non-pathogenic relative, *S. punctatus*. This method lays the foundation for molecular genetic tools needed to establish ecological mitigation strategies and answer broader questions in evolutionary and cell biology.

## Introduction

Basal to the multicellular Dikarya that include both Ascomycota (e.g. sac fungi) and Basidiomycota (e.g. mushrooms) (**Fig. 1A**),^1–3^ Zoosporic fungi have retained many ancestral phenotypes including motile, unicellular, flagellated cell types called zoospores.^4,5^ Crucial for dispersal, zoospores likely rely on complex sensory-motor coordination to locate and settle at nutrient-rich sites.^6–9^ Settled zoospores quickly grow into sporangia, inside which 20-50 new zoospores develop to begin the cycle anew (**Fig. 1B**).

**Figure 1.**
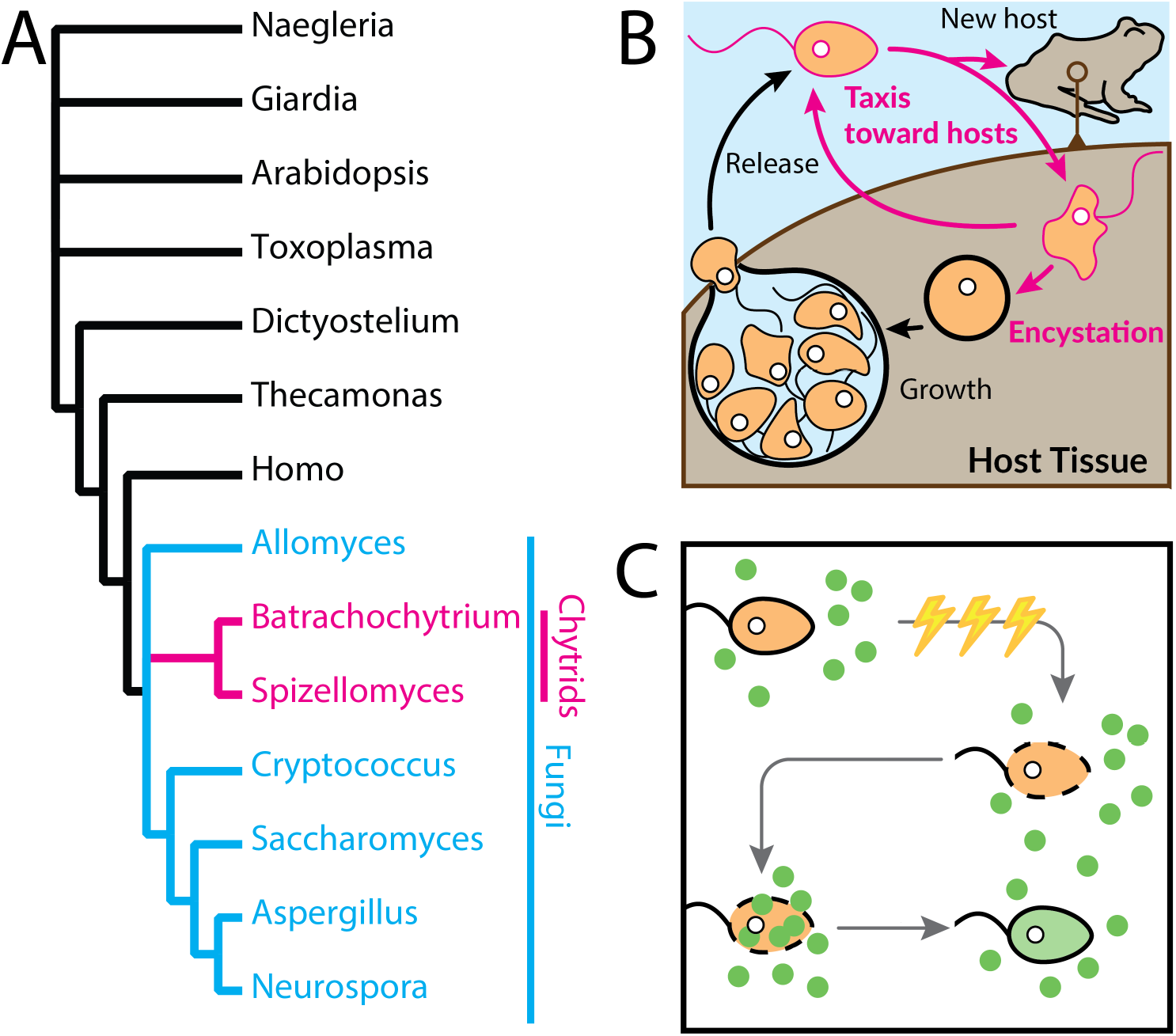
**(A)** phylogenetic tree showing the position of fungi relative to other notable groups across the tree of life. Note the basal position of divergence between Chytrids and all other fungi. **(B)** A diagram showing the major life stages of pathogenic chytrid fungi. Zoospores are released from a sporangia, swim to new hosts or reinfect the old host. They attach, perhaps crawling to a favorable settlement site where they encyst, synthesize a chitinous cell wall, and invade host cells. The encysted zoospore swells in size, undergo rounds of mitosis without cytokinesis, divides into more zoospores, and opens—releasing them to start the cycle anew. **(C)** Simplified diagram showing the process of electroporation. Electric shocks are applied to a cell (orange) in the presence of a fluorescent payload molecule (green). This opens holes in the cell membrane, allowing the payload to invade the cytoplasm. Over time, the cell membrane repairs and the cell continues to develop with the payload inside.

In the last two decades, chytrids in the genus *Batrachochytrium* have emerged as virulent amphibian pathogens and are the first known example of a chytrid-driven epidemic in vertebrates.^10,11^ Within the zoosporic phylum Chytridiomycota (chytrids), *Batrachochytrium dendrobatidis (Bd)* and *B. salamandrivorans* (*Bsal*) infect amphibians, resulting in mortality rates that have driven multiple species extinct.^12–14^ Although exact numbers are currently under analysis, *Bd* alone has led to a catastrophic global decline of amphibians and *Bsal* is now considered an emerging global pathogen.^12,15,16^ Exacerbated by climate change and anthropogenic distribution, it is unclear how much additional amphibian diversity will be lost to chytrid infection without intervention.^17,18^

A significant hurdle to effective management of chytrid infections is the lack of basic knowledge about their biology. With no way to study the molecular mechanisms underlying host identification, infection, and growth in parasitic chytrids, the field is struggling to develop mitigation strategies for even a single host species. While tools are just emerging for non-parasitic relatives^19^, developing molecular genetic tools for parasitic chytrids will remove key roadblocks hindering our efforts to curb the pathogens’ ecological impact. Additionally, these tools will leverage the unique diversity and evolutionary history of zoosporic fungi to better understand broader principles underlying the origins of parasitism, transitions to pathogenicity, and evolutionary cell biology.

Molecular genetic tools generally require driving molecular payloads across the cell membrane. To deliver molecules to the cytoplasm of chytrid cells, we have optimized one such method, electroporation, because of its widespread use in fungi and the flexibility of its downstream applications.^20–22^ Electroporation exposes cells to tightly controlled electrical pulses that create holes in the cell membrane. The now-permeable membrane allows for co-incubated compounds to passively enter the cell before the membrane is sealed (**Fig. 1C**). Successful electroporation depends largely on physical parameters such as cell size, pulse profile, buffer conductance, and electrode contact area,^20,23^ each of which must be optimized for a particular cell type. The comparatively small role that biological parameters play in the success of electroporation make it applicable—given enough persistence—to cells from almost any taxa.

Although the initial process of developing electroporation for a new species is time-intensive, this investment yields a general template that can be quickly expanded to closely related species and optimized for delivery of a wide variety of payloads. Electroporation is widely used to introduce heterologous DNA into cells and perform CRISPR/Cas9-mediated gene editing, although these methods require significant development beyond electroporation. Conversely, other tools such as antisense oligonucleotides, siRNAs, and small molecule inhibitors can be used almost immediately after electroporation has been optimized.^24–27^

To study the biology of infectious chytrids, we have developed a reliable, high-efficiency electroporation protocol for chytrid fungi. We optimized this protocol for two pathogenic chytrids (*B. dendrobatidis*, *B. salamandrivorans*), and a non-pathogenic chytrid (*Spizellomyces punctatus, Sp*). This protocol opens the door for further studies using molecular genetics to address both ongoing and future chytrid epidemics while laying the foundation for comparative studies in these unique and diverse lineages.

## Results

### Voltage for optimal electroporation varies between chytrid species

To track the success of molecular payload delivery during electroporation, we used fluorescein-labelled dextrans (hereafter referred to simply as dextrans) and measured delivery using both microscopy and flow cytometry. To optimize electroporation, one must explore a large parameter space that includes the variables: pulse shape, pulse time, number of pulses, contact distance, contact surface area, buffer conductivity, and pulse voltage. To quickly explore this parameter space, we conducted single-replicate trials looking for parameter combinations that resulted in at least 15% of single cells loaded with dextrans. This wide exploration led us to an unoptimized protocol using two three-millisecond square-wave pulses, five seconds apart yielding 10-20% efficiency (data not shown). To optimize this protocol for *Bd*, we adjusted pulse voltage alone testing 750v, 1000v, and 1250v. Over this range, we found efficiency peaked at 1000v, with ~95% of zoospores loaded with dextrans. We compared this efficiency to controls for autofluorescent shift due to pulse exposure (electroporation without dextran incubation) and dextran ingestion or non-specific binding to cell walls/coats (dextran incubation without electroporation). These comparisons revealed the fluorescence of dextran-incubated, electroporated cells is due to successful dextrans delivery into the cytosol. When translating this optimized protocol, with no adjustments, into *Bsal* and *Sp*, we found the optimal voltages were: *Bsal*, 1250v; *Sp*, 1000v (**Figs. 2-4**).

**Figure 2.**
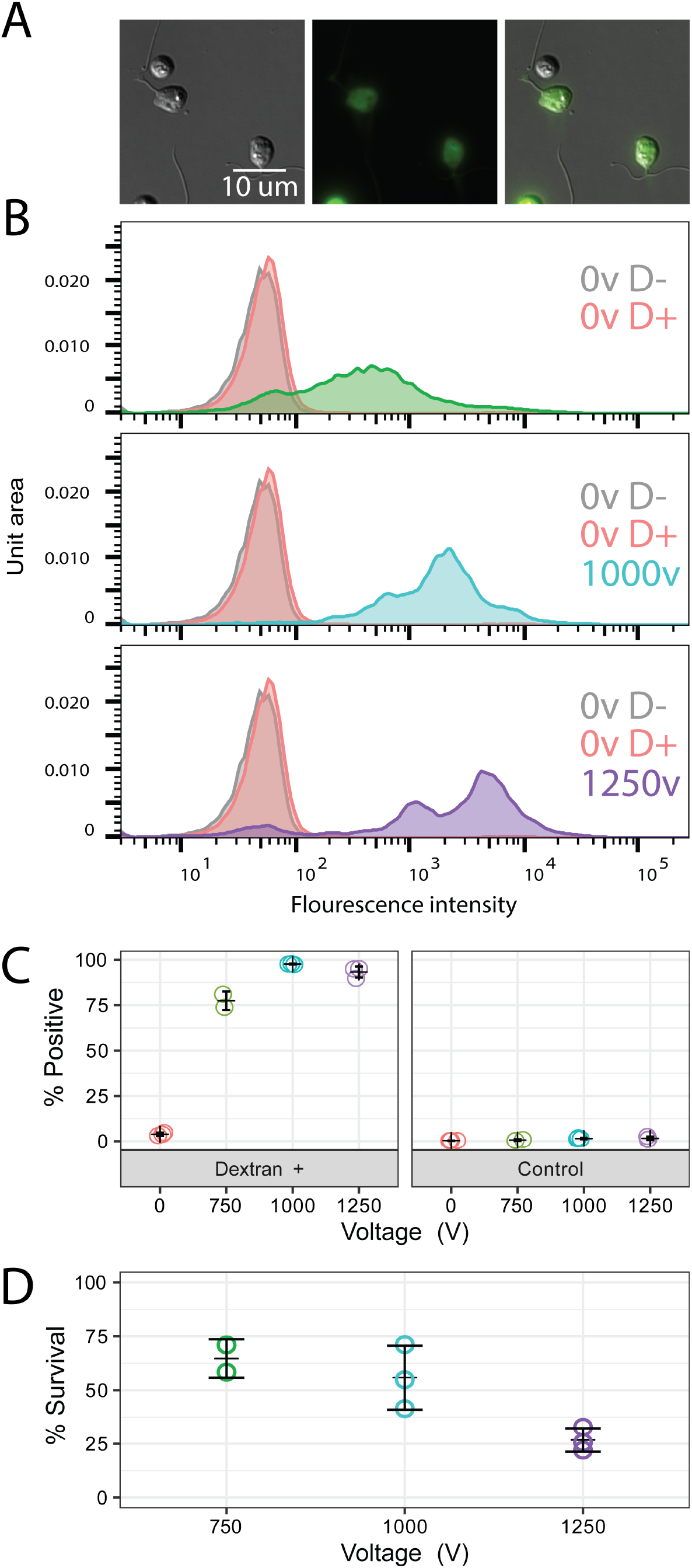
Electroporation efficiency and survival of *B. dendrobatidis* at 750v (**green**), 1000v (**blue**), and 1250v (**purple**). **(A)** Representative images of cells after electroporation at 1000v showing live cells adhered to the slide and attempting amoeboid movement. **(B)** Representative flow cytometry data from a single replicate showing the fluorescence intensity of single cell events for non-electroporated, no dextran controls (*grey*); non-electroporated, dextran incubated controls (*red*); and electroporated, dextrans incubated treatments (*colors vary by voltage*). **(C)** Percent of single cell events with fluorescence intensities above the non-electroporated, no dextran control relative to all single cell events for three independent biological replicates. **(D)** Percent survival of cells in electroporated, dextran incubated controls in three independent biological replicates. Counts were normalized to the respective non-electroporated, no dextran controls.

**Figure 3.**
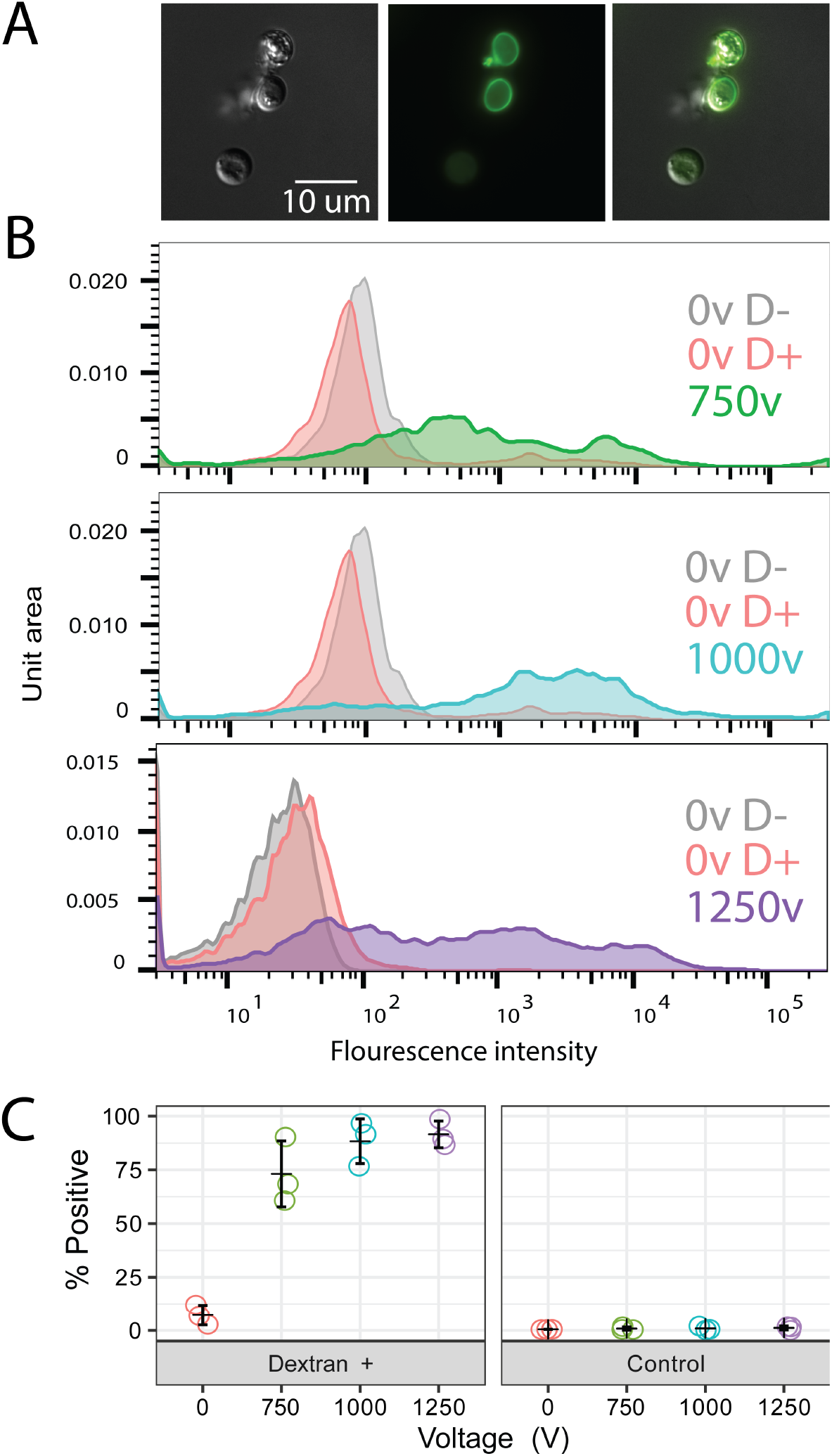
Electroporation efficiency of *B. salamandrivorans* for 750v (**green**), 1000v (**blue**), and 1250v (**purple**). **(A)** Representative images (DIC, FITC, Merged) from a single replicate at 1000v showing positive cells with dextran inside the cell and not only/merely stuck to the cell wall/coat. **(B)** Representative flow cytometry data from a single replicate showing the fluorescence intensity of single cell events for non-electroporated, no dextran controls (*grey*); non-electroporated, dextran incubated controls (*red*); and electroporated, dextran incubated treatments (*colors vary by voltage*). **(C)** Percent of single cell events with fluorescence intensities above the non-electroporated, no dextran control relative to all single cell events for three independent biological replicates.

**Figure 4.**
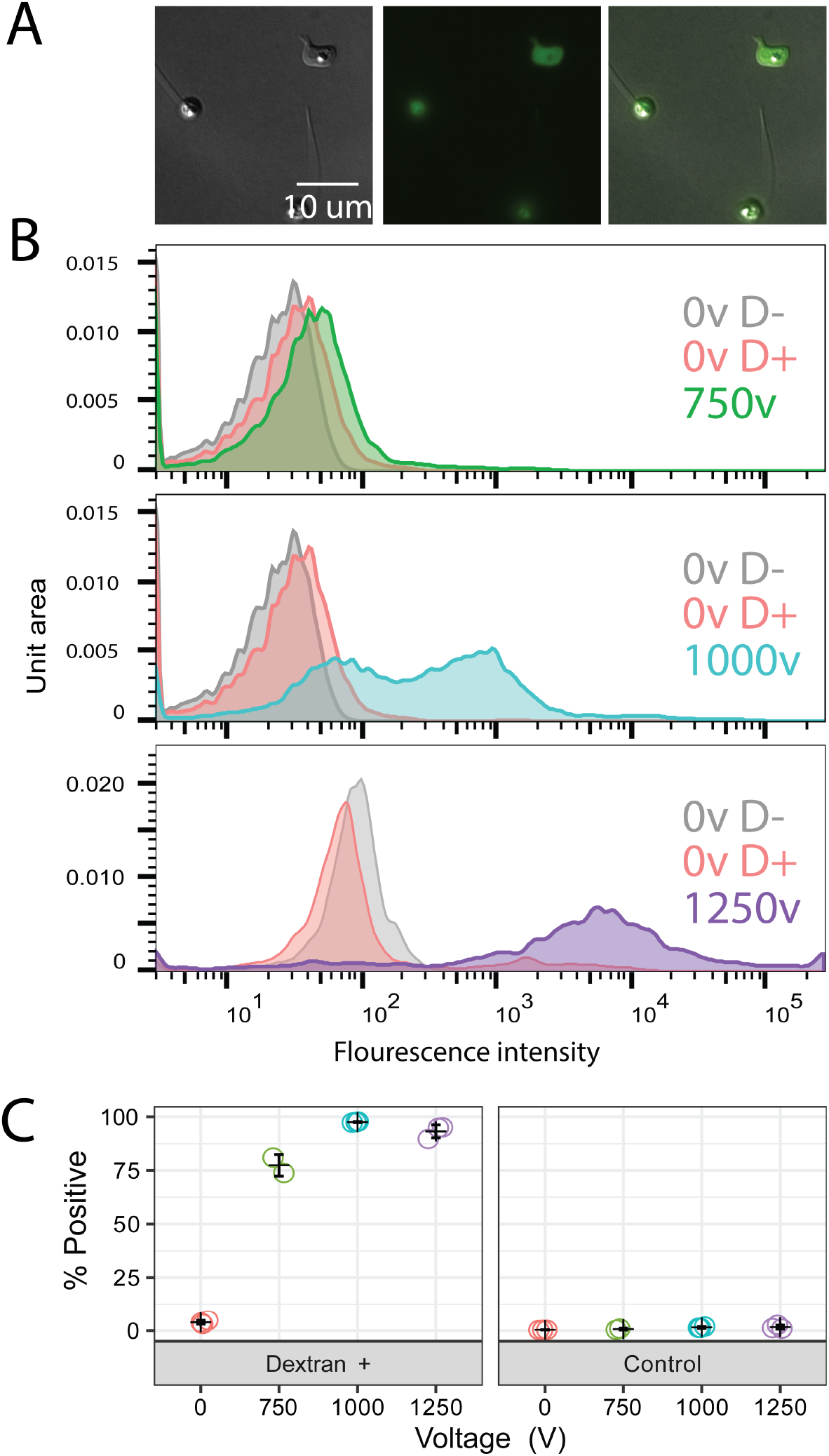
Electroporation efficiency of *S. punctatus* for 750v (**green**), 1000v (**blue**), and 1250v (**purple**). **(A)** Representative cells from a single replicate at 1000v showing positive cells with dextran inside the cell and not stuck to the cell wall/coat. **(B)** Representative flow cytometry data from a single replicate showing the fluorescence intensity of single cell events for non-electroporated, no dextran controls (*grey*); non-electroporated, dextran incubated controls (*red*); and electroporated, dextrans incubated treatments (*colors vary by voltage*). **(C)** Percent of single cell events with fluorescence intensities above the non-electroporated, no dextran control relative to all single cell events for three independent biological replicates.

### Cell walls of encysted spores accumulate fluorescent molecules

When analyzing flow cytometry data, we used non-electroporated, no dextran controls to set a ‘positive’ gate which excluded 99.5% of all control cells. To obtain the percent of positive cells under other treatments, we transposed this gate onto each experimental and control treatment within the experimental replicate. Unexpectedly, non-electroporated spores incubated with dextrans showed 3-10% of cells in the treatment were positive (**Figs. 2-4**). To understand why incubation with dextrans leads to an increase of positive cells in the absence of electroporation, we inspected these samples using fluorescence microscopy. We observed cells with a bright ring of fluorescence on the periphery of a small percentage of cells in all samples incubated with dextrans regardless of their exposure to electric shock (**Fig. 5**), suggesting the small percent of positive cells in non-electroporated samples is due to non-specific binding of dextrans to the cell-coat or cell-wall.

**Figure 5.**
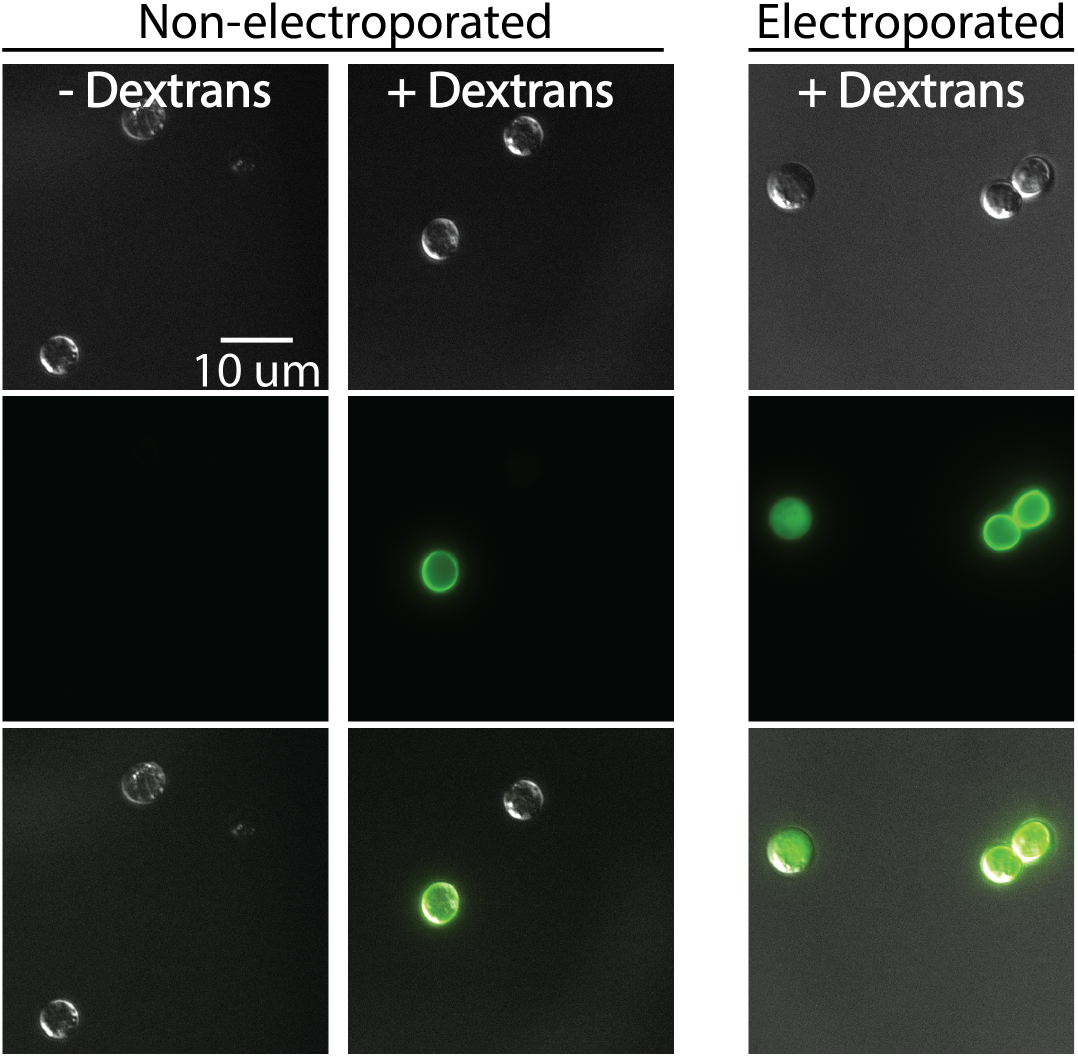
Although dextran staining of cell wall/coat occurs without electroporation, stained and electroporated cells can be distinguished via microscopy. Representative images of each treatment were taken from a single replicate performed on subsets of spores from the same population. Fluorescence intensity was normalized across all images. Non-electroporated cells incubated with dextrans (*non-electroporated, +dextrans*) show weak staining in a pattern that suggests binding to the cell wall. This same pattern can be observed in electroporated samples (*electroporated, +dextrans*) where stained cells (*right*) co-occur with positive cells (*left)* in which dextrans fluorescence is limited to the inside of the cell.

### Electroporation efficiency is strongly affected by dextran manufacturer

To explore if cheaper sources of 3000MW dextrans could substitute our initial choice (Thermo-Fisher), we tested an alternative manufacturer of 3000MW dextrans (Sigma-Aldrich). In the lots and concentrations we tested under recommended storage conditions, we find a 30-70% difference in loading efficiency between two dextrans manufacturers (**Fig.S2**). In side-by-side comparisons, Thermo Fisher anionic 3000MW dextrans (Thermo, D3305) yield 75-97% loading efficiency while the cheaper option (**Fig. S1A**), Sigma 3000MW dextrans (Sigma, FD4) results in a lower and more variable efficiency (**Fig. S1B**). After electroporation, Sigma FD4 dextrans are confined to a small point in the majority of cells but do not stain the cell wall (**Fig. S1B**). In contrast, Thermo-Fisher dextrans appear evenly distributed throughout the cytoplasm in the majority of cells with occasional staining of the cell wall (**Fig. S1A**).

### Synchronizing spores is not required for high efficiency

Without synchronizing cultures, chytrid zoospores are different ages and represent a very heterogeneous population of cells ranging from just released to about to encyst.^28^ To test if zoospore age had an impact on electrocompetency, we synchronized cultures to limit the confounding variables. Once we achieved reproducible and effective electroporation with synchronized cultures, we tested if zoospore synchrony would affect efficacy. We find that although synchronization of zoospores may slightly increase efficiency, it is not required to reach > 80% loading of all cells (**Fig. S2**). Use of synchronized spores, however, may facilitate downstream applications of electroporated zoospores.

## Methods

### Culture and synchronization of zoospores

We cultured *Batrachochytrium* and *Spizellomyces* species in tissue culture treated flasks (Celltreat 229330). Strains were thawed and then routinely passaged according to the following species-specific protocols: We grew *B. dendrobatidis* (JEL423) in 1% Tryptone for 3 days at 24°C *B. salamandrivorans* (AMFP1) in ½ strength TGHL for 4 days at 15°C, and *Spizellomyces punctatus* (ATCC 48900) on K1 plates at 30°C for 1-2 days. After the fungi were mature, we collected zoospores and seeded the next passage.

For *Batrachochytrium* species, we synchronized zoospores by discarding growth media and gently washing older zoospores away from the sporangia adhered to the tissue culture flask three times with fresh growth media. After two hours of incubation with fresh growth media, we collected synchronized zoospores for use. For *Spizellomyces* species, we checked for mature sporangia after 19-32 hours of growth while ensuring the surface of the agar had a thin film of liquid around each sporangium. As soon as sporangia began releasing zoospores, we washed plates three times with 2mL of K1 media and let the sporangia sit under another 2mL for 30 minutes before collecting synced zoospores.

### Electroporation

We have made the detailed electroporation protocol available on protocols.io (https://www.protocols.io/private/974AC160056C11EAA3CF0242AC110004) and in a notebook-ready checklist format (**File S1**). In short, zoospores are brought to a specific concentration, co-incubated on ice with dextrans, electroporated using high-voltage square-wave pulses, then allowed to recover on ice. Unless specifically noted, all solutions were sterilized and we synchronized zoospores prior to collection. To achieve proper buffer conductance during electroporation, we transferred zoospores into SM buffer (5mM KCl, 15mM sodium phosphate buffer pH 7.2, 15mM MgCl_2_, 25mM sodium succinate dibasic hexahydrate, 25mM D-Mannitol) after collection by repeating the following wash steps three times: centrifuge zoospores at for 5min at 2500rcf, remove supernatant, then resuspend in 5mL of SM buffer. We then brought zoospores to a final concentration of 1×10^7^ zoospores/mL. In preparation for electroporation, we loaded 100uL of this 1×10^7^ zoospores/mL suspension into a 0.2cm electroporation cuvette (Bulldog Bio 12358-346) along with either: 100uL of 2mg/mL 3000MWCO fluorescein-labelled dextrans (Thermo-Fisher, Cat. no. 3305) in SM buffer for experimental treatments, or 100uL SM buffer for control treatments. We then chilled all cuvettes on ice for 10 min. Immediately before electroporation, we gently mixed each cuvette by pipetting up and down to resuspend chilled zoospores. To electroporate cells, we used two, three millisecond, square-wave pulses five seconds apart (Biorad GenePulser Xcell + CE module) for all samples. We tested 750v, 1000v, and 1250v for all species. We immediately placed each cuvette on ice for 10 minutes after electroporation then gently added 200uL of 1% tryptone and allowed cells to rest for another 10 minutes on ice.

### Cytometry and Microscopy measurement of electroporation efficiency

We measured parameter efficiency using flow cytometry and used microscopy to verify that cells were loaded, rather than just externally stained. These steps are not necessary to grow electroporated cells for downstream applications and can be applied to a subset of cells, or even skipped after optimizing the protocol for new payloads. To prepare cells for flow and microscopy, we transferred all 400uL from the electroporation cuvette into 5mL of chilled media. To remove extracellular dextrans and autofluorescent media, we performed a series of 6 wash steps. For each step, we centrifuged zoospores at 2500 rpm for 5 minutes at then discarded the supernatant and gently resuspended the zoospore pellet in 5 mL of the indicated solution. For the first 3 wash steps, we resuspended zoospores in chilled growth media; for the next 2 wash steps we resuspended in room-temperature SM buffer. For the final wash step in SM buffer, we resuspended the zoospores using only 600uL of SM buffer. We then transferred 100uL of zoospores to a separate tube for imaging and fixed the remaining volume by adding an equal volume of paraformaldehyde (PFA) based fixation buffer, final concentrations: 4.8% PFA, 9 mM sucrose, 50mM Sodium Phosphate Buffer pH 7.2. We incubated the zoospores and fixation buffer on ice for 20 minutes, after which we centrifuged them for 5 minutes at 2200 rpm, discarded the supernatant, and resuspended in 400uL of unchilled SM buffer.

We used flow cytometry of the fixed cells to measure the overall ratio of loaded to unloaded zoospores. Using a BD LSR Fortessa 3 Laser cytometer and BD FACSDiva software, we captured 10,000 events and analyzed the resulting data using FlowJo v10.1. Cells were gated for enhanced FITC in single-cell events relative to control treatments. To corroborate flow cytometry measurements, we imaged live cells on a Nikon Ti2-E inverted microscope equipped with a 100x PlanApo objective and sCMOS 4mp camera (PCO Panda) using NIS Elements software to capture images in FITC (LED illuminator at 488nm) and DIC (white LED transmitted light) images. *Bd* zoospores were adhered to the slide by pre-treating it with concanavalin-A,^29^ all other spores were imaged without treating the slide.

## DISCUSSION

Developing efficient, reproducible molecular tools for new species remains a crucial step towards unravelling the general principles, origins, and evolution of basic cell biology. We present a well-optimized electroporation protocol for the zoospores of infectious chytrid fungi, with between 75-97% of cells demonstrably loaded with a fluorescent dextran payload. The efficiencies we report are likely conservative because electroporated cells are remarkably fragile and the stringent wash steps we used to quantify the percent of cells loaded likely resulted in additional cell death that could be avoided in typical applications. With reliable electroporation now established, the field can begin developing refined molecular genetic tools, such as CRISPR, to empirically test hypotheses about the molecular mechanisms underlying chytrid virulence and zoospore evolution.

### Potential pitfalls and difficulties

Despite the general convenience of electroporation, we find that there are a number of qualitative variables which can have a large effect on the replicability of this protocol. For instance: using a cheaper source of what should be identical dextrans results in a 30-70% drop in efficacy but using asynchronized cells produces no noticeable effect on efficacy. In an effort to help future researchers, we discuss a number of pitfalls and difficulties we believe will help make applications of this protocol successful:

#### Zoospores are fragile

We observed a large drop in efficacy and single cell events measured by flow cytometry if cells were pipetted or poured too roughly at any stage of the protocol. Thus, the use of any shaker/vortexer is not recommended at any point in this protocol before cell fixation. Researchers attempting this protocol should also exercise caution and be sure to pipette gently when resuspending spores. We also noted a qualitative increase in cell death at the end of this protocol if cells experienced large temperature fluctuations and advise that cuvettes are initially loaded with room-temperature reagents, then everything be placed and kept on ice after the cuvettes have been loaded to allow the cells to chill gradually.

#### Cell walls bind dextrans

Qualitative image analysis suggests that “stained” cells all lack a visible flagellum and therefore have likely begun the encystment process. This process is also marked by rapid synthesis of cell wall, endocytosis, and growth.^30,31^ We find it likely that dextrans in the external environment bind to the newly synthesized chitin cell wall similar to the binding of wheat germ agglutinin to saccharides in fungal cell walls.^32^ Without confirmation and the establishment of methods to quench externally bound FITC molecules, the use of “no electroporation” controls incubated with dextrans are necessary to estimate the percentage of cells which are ‘positive’ but do not have dextrans in the cytoplasm.

#### Loss of fluorescence in fixed cells after 6 hours

After approximately 6 hours, dextran loaded cells were not detectable by flow cytometry, even when re-sampling previously fluorescent samples stored in the dark at 4°C overnight. This is likely a result of using non-fixable fluorescein. If users elect to use the same dextran as we did in these experiments (D3305) we urge them to avoid permeabilization steps and to proceed as quickly as possible from fixation to flow cytometry. Alternatively, users may use Thermo Cat. No. **D3306**, a lysine-fixable 3000MW fluorescein labelled dextran if they would like to leave fixed samples overnight or include additional stains that require permeabilization.

#### Zoospore viability after electroporation

Most downstream applications of this protocol will require cells to remain alive after electroporation. We confirmed that spores loaded with dextrans were alive, active, and crawling at least 4 hours after electroporation. However, the survival rate of electroporated spores is likely to be heavily dependent on the molecules introduced into the cell. While not necessary to assess the efficacy of electroporation itself, any future application of this protocol should assess the halflife of their introduced molecule through flow cytometry, interrupted phenotypes, or antibiotic resistance.

## Conclusions

Developing molecular tools for chytrids allows researchers access to unique, diverse lineages for molecular studies into the origin, evolution, and fundamental processes underlying cell biology and parasitism. Our protocol is the first step towards building a suite of molecular tools allowing us to study chytrid fungi in depth. We anticipate this protocol will give rise to a multitude of studies using morpholinos, small molecule inhibitors, siRNAs, and eventually DNA editing to investigate the mechanisms and evolution behind chytrid biology. Furthemore, our results suggest that the protocol presented here is likely universal within Chytridiomycota and may be used as a starting template for other zoosporic species. Zoosporic fungi are now poised to quickly advance solutions to current ecological problems and provide context for broader questions in evolutionary and sensory cell biology.

## Acknowledgements

This material is based upon work supported by the National Science Foundation under Grant No. IOS-1827257 (LKF-L).

## Competing interests

The authors declare no competing interests.

## Author contributions

All authors contributed to experimental design and performed experiments. A.S. and L.F-L. produced the manuscript.

## Notes

### Competing Interest Statement

The authors have declared no competing interest.

## References

1. James, T. Y. et al. Reconstructing the early evolution of Fungi using a six-gene phylogeny. Nature 443, 818–822 (2006).

2. James, T. Y. et al. A molecular phylogeny of the flagellated fungi (Chytridiomycota) and description of a new phylum (Blastocladiomycota). Mycologia 98, 860–871 (2006).

3. Powell, M. J. & Letcher, P. M. 6 Chytridiomycota, Monoblepharidomycota, and Neocallimastigomycota. in Systematics and Evolution 141–175 (Springer, 2014).

4. James, T. Y., Porter, T. M. & Wallace Martin, W. Blastocladiomycota. in Systematics and Evolution 177–207 (Springer Berlin Heidelberg, 2014).

5. Fuller, M. S. The Zoospore, Hallmark of the Aquatic Fungi. Mycologia 69, 1–20 (1977).

6. Avelar, G. M. et al. A rhodopsin-guanylyl cyclase gene fusion functions in visual perception in a fungus. Curr. Biol. 24, 1234–1240 (2014).

7. Machlis, L. Zoospore Chemotaxis in the Watermold Allomyces. Physiol. Plant. 22, 126–139 (1969).

8. Saranak, J. & Foster, K. Rhodopsin guides fungal phototaxis. Nature 387, 465–466 (1997).

9. Swafford, A. J. M. & Oakley, T. H. Multimodal sensorimotor system in unicellular zoospores of a fungus. J. Exp. Biol. 221, (2018).

10. Longcore, J. E., Pessier, A. P. & Nichols, D. K. Batrachochytrium Dendrobatidis gen. et sp. nov., a Chytrid Pathogenic to Amphibians. Mycologia 91, 219–227 (1999).

11. Fisher, M. C. & Garner, T. W. J. Chytrid fungi and global amphibian declines. Nat. Rev. Microbiol. (2020) doi:10.1038/s41579-020-0335-x.

12. Scheele, B. C. et al. Amphibian fungal panzootic causes catastrophic and ongoing loss of biodiversity. Science 363, 1459–1463 (2019).

13. Van Rooij, P. et al. Germ tube mediated invasion of Batrachochytrium dendrobatidis in amphibian skin is host dependent. PLoS One 7, e41481 (2012).

14. Moss, A. S., Reddy, N. S., Dortaj, I. M. & San Francisco, M. J. Chemotaxis of the amphibian pathogen Batrachochytrium dendrobatidis and its response to a variety of attractants. Mycologia 100, 1–5 (2008).

15. Lambert, M. R., Womack, M. C. & Byrne, A. Q. Comment on ‘Amphibian fungal panzootic causes catastrophic and ongoing loss of biodiversity’. (2020).

16. Scheele, B. C. et al. Response to Comment on ‘Amphibian fungal panzootic causes catastrophic and ongoing loss of biodiversity’. Science vol. 367 (2020).

17. Cohen, J. M., Civitello, D. J., Venesky, M. D., McMahon, T. A. & Rohr, J. R. An interaction between climate change and infectious disease drove widespread amphibian declines. Glob. Chang. Biol. 25, 927–937 (2019).

18. Louca, S., Lampo, M. & Doebeli, M. Assessing host extinction risk following exposure to Batrachochytrium dendrobatidis. Proc. Biol. Sci. 281, 20132783 (2014).

19. Medina, E. M. et al. Genetic transformation and live-cell nuclear and actin dynamics during the life cycle of a chytrid. bioRxiv 787945 (2019) doi:10.1101/787945.

20. Thompson, J. R., Register, E., Curotto, J., Kurtz, M. & Kelly, R. An improved protocol for the preparation of yeast cells for transformation by electroporation. Yeast 14, 565–571 (1998).

21. Chakraborty, B. N., Patterson, N. A. & Kapoor, M. An electroporation-based system for high-efficiency transformation of germinated conidia of filamentous fungi. Can. J. Microbiol. 37, 858–863 (1991).

22. Chakraborty, B. N. & Kapoor, M. Transformation of filamentous fungi by electroporation. Nucleic Acids Res. 18, 6737 (1990).

23. Hui, S. W. Effects of Pulse Length and Strength on Electroporation Efficiency. in Plant Cell Electroporation and Electrofusion Protocols (ed. Nickoloff, J. A.) 29–40 (Springer New York, 1995).

24. Xu, X. et al. Efficient homology-directed gene editing by CRISPR/Cas9 in human stem and primary cells using tube electroporation. Sci. Rep. 8, 11649 (2018).

25. Kotnik, T. et al. Electroporation-based applications in biotechnology. Trends Biotechnol. 33, 480–488 (2015).

26. Fan, Y. & Lin, X. Multiple Applications of a Transient CRISPR-Cas9 Coupled with Electroporation (TRACE) System in the Cryptococcus neoformans Species Complex. Genetics 208, 1357–1372 (2018).

27. Dev, S. B., Rabussay, D. P., Widera, G. & Hofmann, G. A. Medical applications of electroporation. IEEE Trans. Plasma Sci. IEEE Nucl. Plasma Sci. Soc. 28, 206–223 (2000).

28. Kristyn A. Robinson, Kenzie E. Pereira, Molly C. Bletz, Edward Davis Carter, Matthew J. Gray, Jonah Piovia-Scott, John M. Romansic, Douglas C. Woodhams, Lillian Fritz-Laylin. Isolation and maintenance of Batrachochytrium salamandrivorans cultures. Dis. Aquat. Organ. doi:10.3354/dao03488.

29. Fritz-Laylin, L. K., Lord, S. J. & Mullins, R. D. WASP and SCAR are evolutionarily conserved in actin-filled pseudopod-based motility. J. Cell Biol. 216, 1673–1688 (2017).

30. Held, A. A. Encystment and germination of the parasitic chytrid Rozella allomycis on host hyphae. Can. J. Bot. 51, 1825–1835 (1973).

31. Koch, W. J. Studies of the Motile Cells of Chytrids. 6. The Monoblepharidales and Blastocladiales Types of Posteriorly Uniflagellate Motile Cell. Mycologia 61, 422–426 (1969).

32. Privat, J. P., Delmotte, F., Mialonier, G., Bouchard, P. & Monsigny, M. Fluorescence studies of saccharide binding to wheat-germ agglutinin (lectin). Eur. J. Biochem. 47, 5–14 (1974).

