## Supplemental Figure 1 for "High-Efficiency Electroporation of Chytrid Fungi"

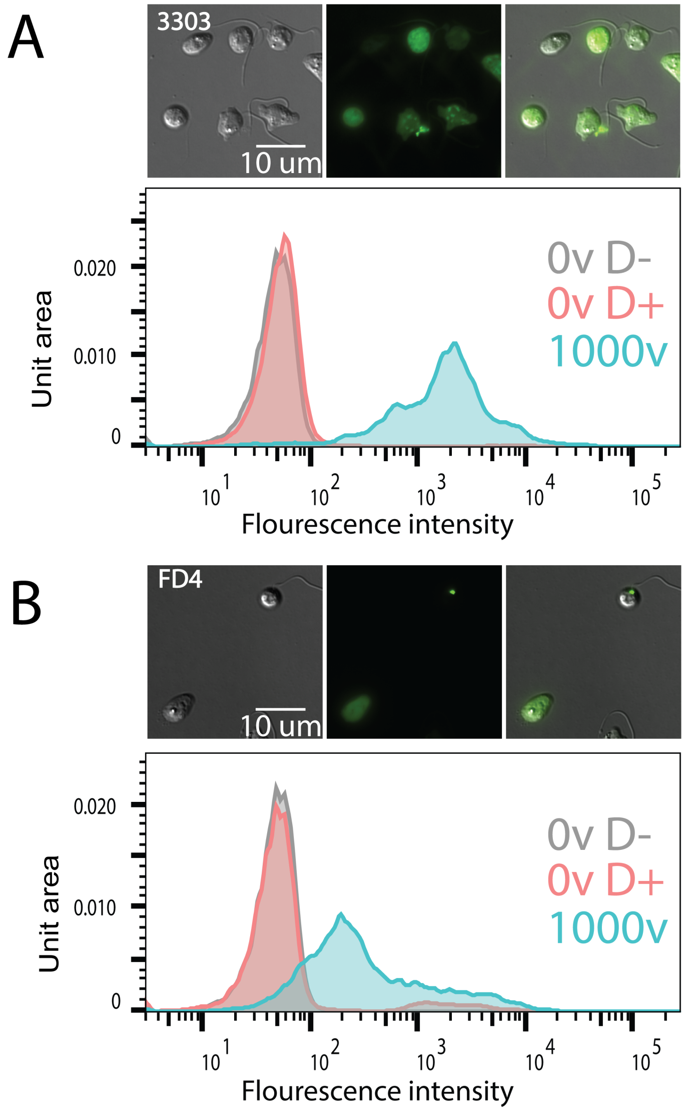


**Supplemental Figure 1**. This protocol is optimized for a single dextran manufacturer. **A)** Representative images and flow cytometry data for a single replicate using Thermo-Fisher Invitrogen dextran (Cat. no. 3303), the manufacturer used throughout the rest of this paper. Clearly positive cells can be observed via microscopy with dextran distributed evenly throughout the cell. **B)** Representative images and flow cytometry data for a single replicate in the same experiment as **(A)** using Sigma dextran (Cat. no. FD4). While positive cells can be observed by microscopy (bottom cell), the majority of cells appear with dextran staining restricted to a small point within the cell as seen in the top cell. Difference is also observed in the flow cytometry data, showing the 1000v, D+ experimental population with left-shifted fluorescence intensity relative to the Invitrogen 3303 dextran.
