## Supplemental Figure 2 for "High-Efficiency Electroporation of Chytrid Fungi"

A

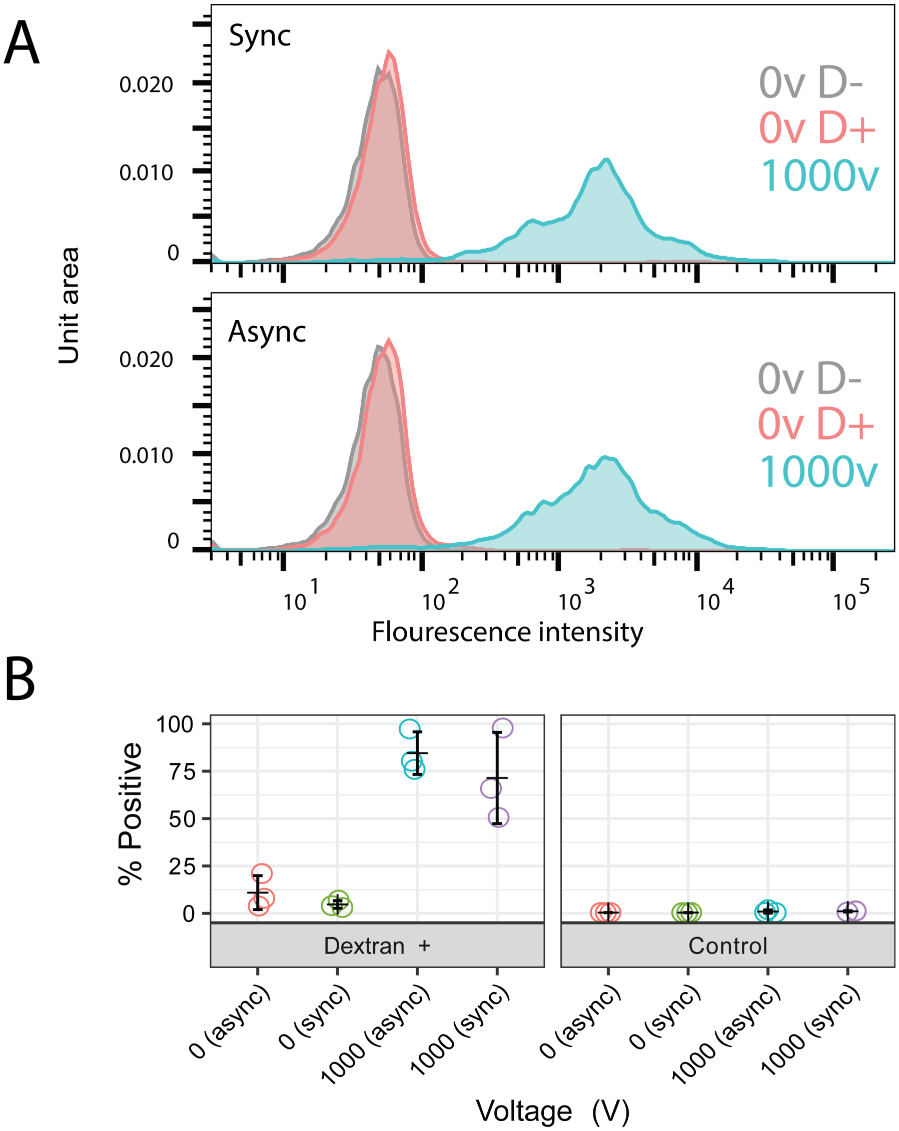


**Supplemental Figure 2**. Synchronizing spores does not grossly affect protocol efficiency. **(A)** Representative flow cytometry data from a single replicate showing the fluorescence intensity of single cell events for non-electroporated, no dextran controls (*grey*); non-electroporated, dextran incubated controls (*red*); and electroporated, dextrans incubated 1000v treatments (*teal*) for synchronized (sync) and non-synchronized (async) cells. **(B)** Percent of single cell events with fluorescence intensities above the non-electroporated, no dextran control relative to all single cell events for three biological replicates.
